# Perceptual axioms are irreconcilable with Euclidean geometry

**DOI:** 10.1101/2023.07.31.551301

**Authors:** Semir Zeki, Zachary F Hale, Ahmad Beyh, Samuel E Rasche

## Abstract

There are different definitions of axioms, but the one that seems to have general approval is that axioms are statements whose truths are universally accepted but cannot be proven; they are the foundation from which further propositional truths are derived. Previous attempts, led by David Hilbert, to show that all of mathematics can be built into an axiomatic system that is complete and consistent failed when Kurt Gödel proved that there will always be statements which are known to be true but can never be proven within the same axiomatic system. But Gödel and his followers took no account of brain mechanisms that generate and mediate logic. In this largely theoretical paper, but backed by previous experiments and our new ones reported below, we show that in the case of so-called “optical illusions” there exists a significant and irreconcilable difference between their visual perception and their description according to Euclidean geometry; when participants are asked to adjust, from an initial randomised state, the perceptual geometric axioms to conform to the Euclidean description, the two never match, although the degree of mismatch varies between individuals. These results provide evidence that perceptual axioms, or statements known to be perceptually true, cannot be described mathematically. Thus the logic of the visual perceptual system is irreconcilable with the cognitive (mathematical) system and cannot be updated even when knowledge of the difference between the two is available. Hence no one brain reality is more “objective” than any other.

## INTRODUCTION

A set of axioms from which geometric truths arise are those of Euclidean geometry, as laid out in Euclid’s *Elements* (Casey, 1885), which constituted the first systematic approach to geometry. This system consists of postulates of planar geometry from which new propositions to describe the physical world can be logically derived. For example, one such axiom states that “things which are equal to the same are equal to one another” (see Figure 1 for more examples). It was thought to be the only system leading to accurate and correct geometrical knowledge of physical space until the 19th century, when new geometries were developed (Gray & Ferreirós, 2022).

**Figure 1.**
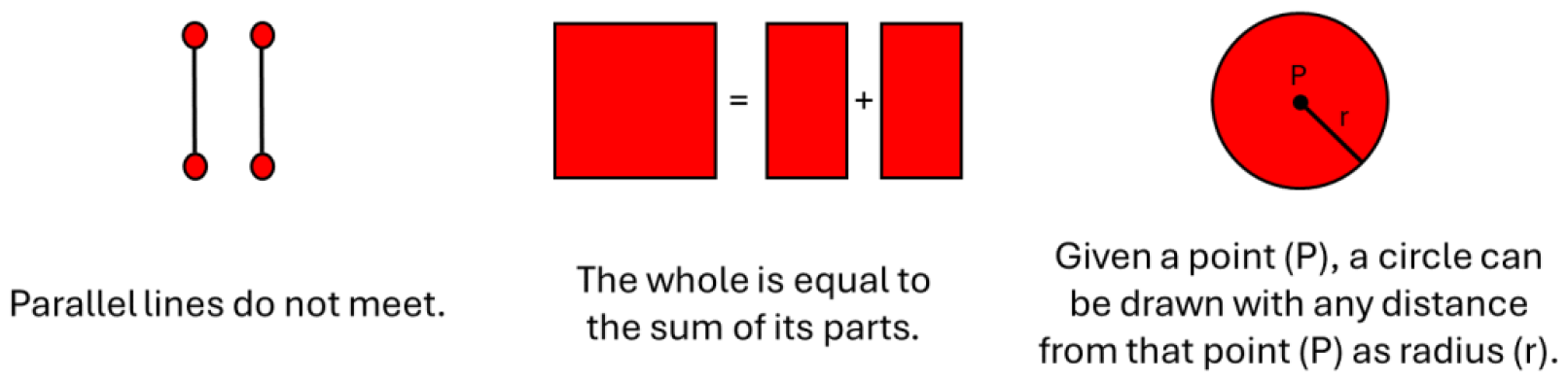
Euclidean Axioms. Examples of postulates/axioms described in Euclid’s *Elements* (Casey, 1885).

Statements of mathematical truths are often considered superior to knowledge acquired by different means, such as perceptual processes; the latter, apparently departing from rigid mathematical precision, are therefore commonly regarded as “illusions”. It was posited recently that there exist alternative truthful axiomatic statements about the organization of our world (Filippov & Zeki, 2022); these are determined by the organization of the visual brain and its logic, which differs from that dictating mathematical axioms; it is irreconcilable with the mathematical system though no less real and compelling. We refer to this system as the perceptual axiomatic system in the sense that, just like the mathematical axiomatic systems, it leads to universally accepted truths that cannot be proven (Filippov & Zeki, 2022). Comparison of perceptual axioms with geometric axioms leads us to conclude that the brain has more than one logical system to dictate the nature of knowledge that it acquires about the world, and the conclusions derived from these different brain logical systems are not necessarily reconcilable. In the two systems we are considering here – the perceptual axiomatic system and the Euclidean geometric one – neither is more objective than the other, because they are both objective, though in different and irreconcilable ways. By axiomatising the statements of perceptual truths, then, we are setting a precedent by which axioms outside the (cognitive) logical deductive system of the brain might challenge the idea that conventional (mathematical and geometric) domains of knowledge are complete.

In this theoretical work, supplemented by experiments, we consider certain unique configurations of two-dimensional geometries, so-called “optical illusions”, where a mathematical description of the geometric components is irreconcilable with the perceptual appearance of those same components. Some of the experiments we report here are similar to experiments done previously (Holt-Hansen, 1961; Knol et al., 2015; Mamassian & de Montalembert, 2010; Prinzmetal et al., 2001; Restle & Decker, 1977; Roberts et al., 2005, *inter alia*), except that the previous experiments were undertaken mainly to determine the parameters affecting the illusory effects. We address a different question here; our aim is to enquire into the extent to which “illusions” depart from the geometric axioms, i.e., to determine whether the two axiomatic systems are reconcilable or not. It is in light of these experiments that we draw a new conclusion, namely that the perceptual axioms are irreconcilable with the Euclidean geometric ones. Importantly, this discrepancy is persistent and self-evident; even when one is informed of the mathematical truth, perception is highly resistant to change.

## METHODS

### Participants

We recruited 28 participants (18 female, 10 male, ages 25.7 ± 5.6) through an online recruitment agency (https://www.callforparticipants.com/) and through internal faculty bulletins. Our pool included participants from nine different ethnic backgrounds, and all had normal or corrected-to-normal vision; all provided informed consent and the experiment was approved by the university’s Research Ethics Committee. The study was conducted at the psychophysics room of the Laboratory of Neurobiology at University College London.

### Stimuli

We selected five geometric optical illusions for their simplicity and ease of (incremental) adjustment. These were the Müller-Lyer, the vertical-horizontal, the Ponzo, the Hering, and the Ebbinghaus illusions (Figure 2). In the Müller-Lyer and Ponzo illusions the lower of two horizontal lines appears shorter than the higher one, whereas in the vertical-horizontal illusion the vertical line appears longer than the horizontal one, when the pairs of lines are mathematically equal. In the Hering illusion the two vertical lines appear convexly curved when they are mathematically parallel. In the Ebbinghaus illusion the central disk surrounded by smaller disks appears larger than the one surrounded by larger disks, when they are mathematically identical.

**Figure 2.**
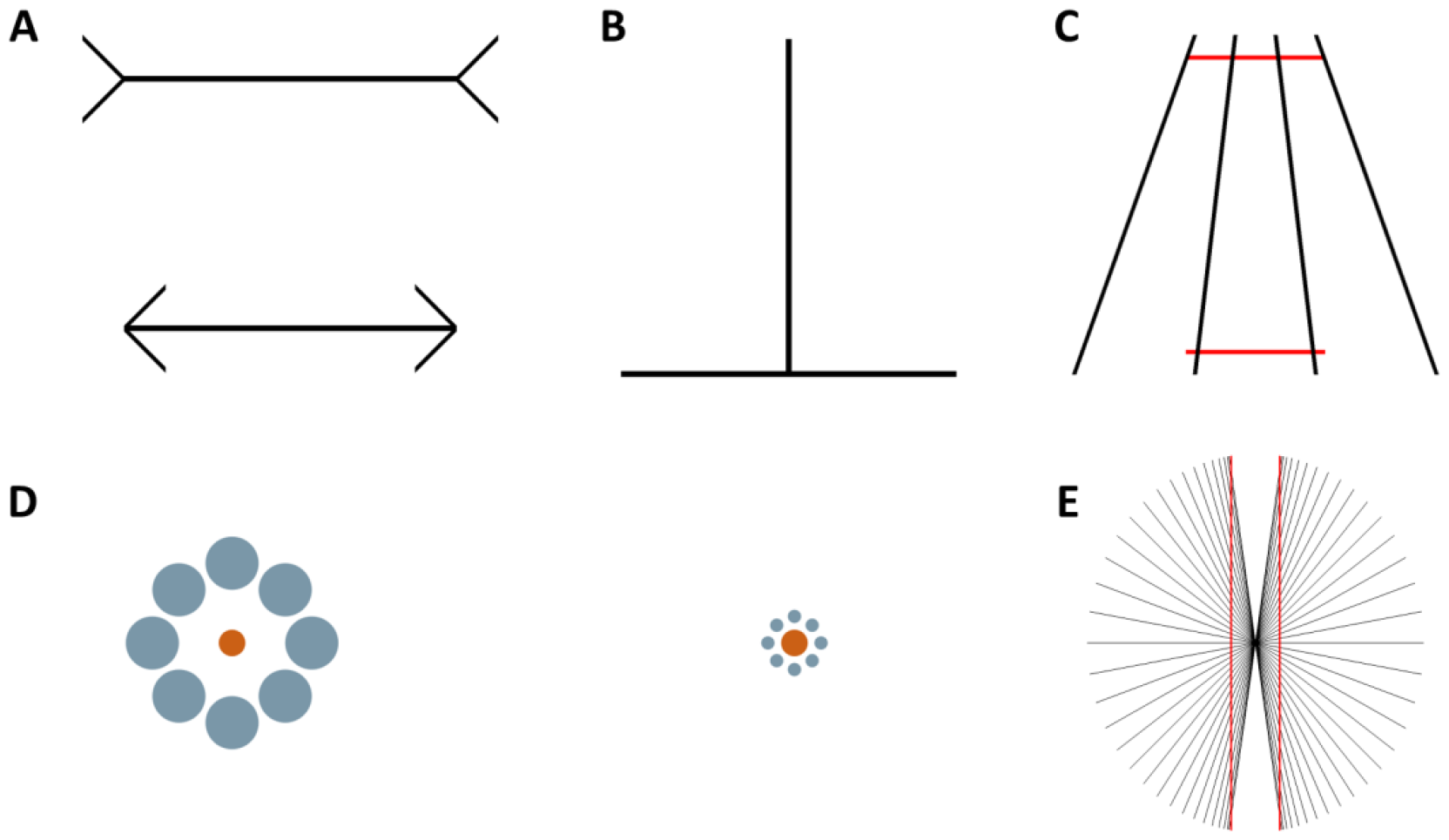
Illusions used in this experiment. A) The Müller-Lyer illusion. B) The vertical-horizontal illusion. C) The Ponzo illusion. D) The Ebbinghaus illusion. E) The Hering illusion.

The illusions used were programmed in Matlab (The Mathworks) using PsychoPhysics Toolbox Version 3 (Brainard, 1997) such that one or more components of each illusion could be adjusted by a fixed increment (or decrement). Each illusion had three starting positions: mathematically truthful (equality), two increments smaller, or two increments larger than the mathematical equality point.

#### Adjustment of the stimuli

When the starting positions were mathematically equal, line lengths in the Müller-Lyer and vertical-horizontal illusions covered 15.9 visual degrees, and the Ponzo illusion 9.2 degrees. Each incremental step, which was carried out by a key press, was equivalent to 0.2 visual degrees on each side of the bottom line for the Müller-Lyer, the Ponzo, and the vertical-horizontal illusions (Figure 2). For the Ebbinghaus illusion, the diameter of the central circles was one visual degree when mathematically equal. The increment of the left inner circle corresponded to a change in diameter of 0.2 visual degrees.

Participants were able to alter the Hering illusion by increasing the curvature of the two lines inwards (toward the central point). This was achieved by modelling the lines with a quadratic formula, x = ay^2^, whereby changing the value of ‘a’ led to a change in the line’s curvature: a = 0 corresponded to a straight line, and a = a_max_ corresponded to the configuration of maximum allowed curvature. The increment corresponded to a = 1; the maximum increment for this illusion was set to a = 30, which corresponded to a range of 1.3 visual degrees from the starting point (straight line) to the peak of the curve.

All stimuli were displayed on a monitor with a resolution of 1920×1080 (W×H) pixels against a plain white background and subtended a maximum of 16° at the eye, from a viewing distance of 60 cm. Pilot data confirmed that the adjustment effected, as a fraction of the absolute size of the initial configuration, was scale-independent (see supplementary material).

### Procedure

Subjects who took part in the study did so in person at the psychophysics room at University College London. The experiment began with a set of preliminary questions where participants indicated whether they perceived the effect of each separate illusion; this amounted to asking whether the two respective lines or circles appeared equal and/or parallel when the shapes are mathematically parallel or equal in length. Participants were told to base their judgements solely on visual perception, rather than prior knowledge or what they thought was veridical. Those illusions for which the participant reported perceiving a geometric asymmetry were then displayed in the task. If a participant indicated not perceiving the illusion, the illusion was not displayed further.

Each illusion was presented randomly five times for 10 s at one of the three initial positions (mathematical truthful, two increments smaller than the mathematically truthful one, or two increments larger), amounting to fifteen presentations in total. During the presentation time, participants had the opportunity to alter the illusion; they were asked to keep adjusting the size of the bottom part of the Müller-Lyer illusion, the Ponzo illusion, and the vertical-horizontal illusion until they appeared equal in length. For the Ebbinghaus illusion, participants were asked to match the size of the two inner disks by adjusting the size of the left inner disk. Adjusting the Hering illusion entailed altering the curvature of the two vertical lines from a state where they appeared to be convex to a state where they appeared to be parallel. Left hand key presses on the keyboard made the lines more concave or smaller, while right hand keypresses made them more convex or larger. If participants were satisfied with the adjustment before 10 s elapsed, they moved on to the next trial by pressing a key. If no response was given during the 10 s period, the trial was terminated and a new one started. However, this 10 s duration was more than enough since participants responded on every trial. A fixation cross was shown for half a second in the interval between rounds, and participants were given a break halfway through the set of trials.

## RESULTS

### Participants’ adjustments

Participants viewed the above five geometric optical “illusions” and were asked to adjust their components from an initial configuration to a final one in which the features appeared equal (see Methods). We began each participant’s testing session by showing them the various stimuli and asking whether they perceived the illusions. Table 1 shows that most participants did, but not all illusions were equally effective, although the magnitude of the modification required to bring the perceptual reality to match the mathematical description (e.g., to make the two lines of the Ponzo illusion appear equal in length) was significant for all illusions.

**Table 1.** Susceptibility to visual illusions and magnitude of the illusory effect.

| Illusion | Proportion of participants who perceived the illusion | Description of modification required to match the mathematical and perceptual equalities | Magnitude of modification | Average individual standard deviation |
| --- | --- | --- | --- | --- |
| Müller-Lyer | 75% | increase length of lower line | 16.1% | 4.8% |
| Vertical-horizontal | 93% | increase length of lower line | 16.6% | 6.1% |
| Ponzo | 93% | increase length of lower line | 48.6% | 7.6% |
| Ebbinghaus | 96% | increase diameter of left disk | 37.4% | 10.6% |
| Hering | 57% | increase curvature of vertical lines | 28.4% | 11.3% |

Most participants (N = 28) were susceptible to the illusory effect of the chosen stimuli, but the stimuli were not all equally effective and required different amounts of modification to transition from mathematical equality to perceptual equality.

Although the magnitude of change varied between participants for any given illusion, it remained relatively stable within each (Table 1, Figure 3). That is to say, all those who perceived the illusion modified the length of it, and did so to a similar degree repetitively, but different subjects modified it to a different extent.

**Figure 3.**
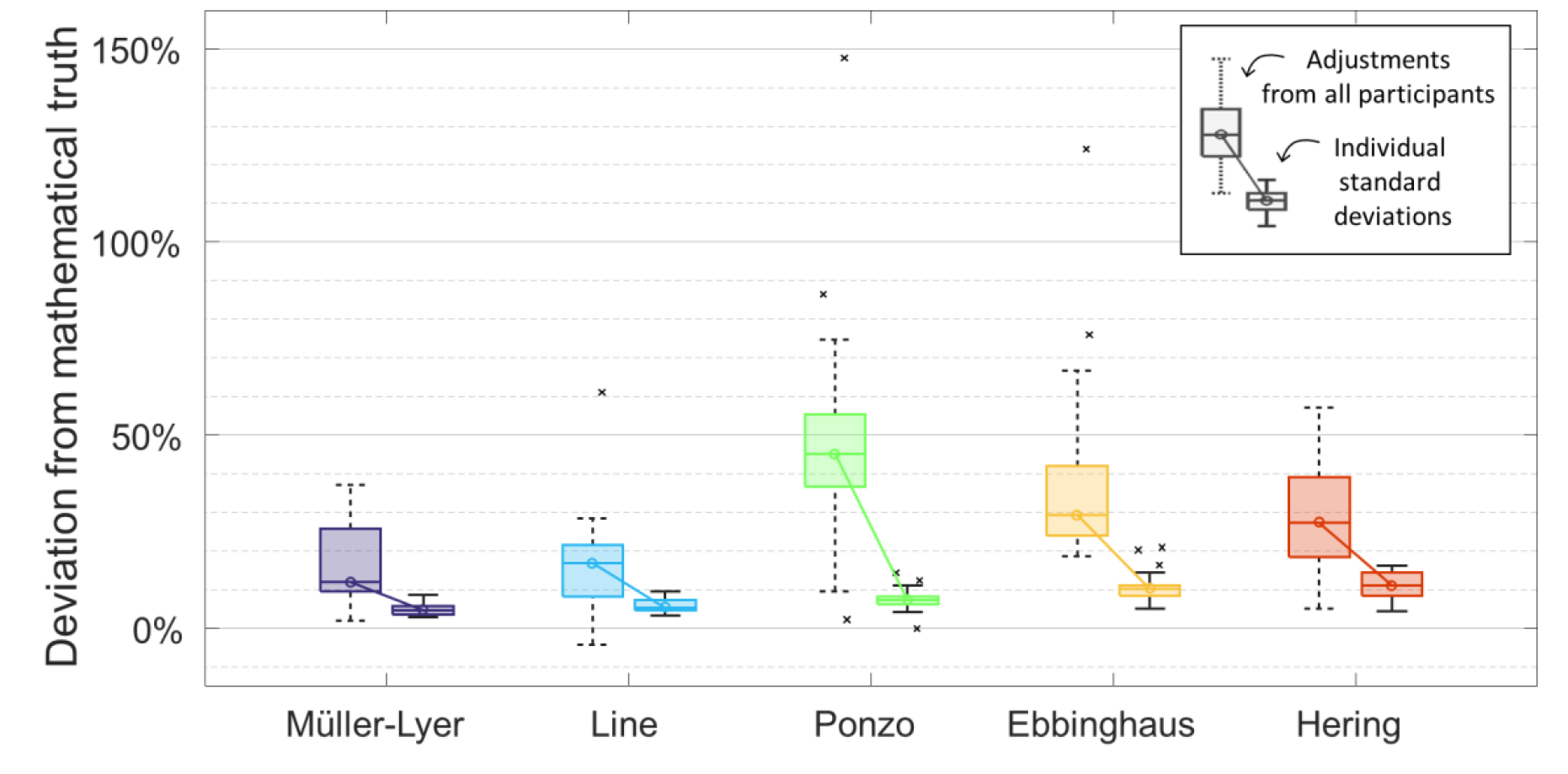
Percentage change from mathematical equality per illusion and standard deviation within participant. For each illusion, the boxplot on the left shows the distribution of the participants’ mean modification in percentages of the original reference size. The boxplot of the same colour on the right shows the distribution of participants’ standard deviations. The left boxplot therefore represents the variability of mean adjustments at the population level, while the right boxplot represents their variability at the individual level. Boxes cover points between the 25^th^ and 75^th^ percentile and central lines and circles indicate the median point of each dataset. The whiskers extend to 1.5x the interquartile range (IQR) from the top/bottom of each box.

## DISCUSSION

In this study, based on the distinction previously made between mathematical and perceptual axioms (Filippov & Zeki, 2022), we sought to demonstrate the irreconcilability between the two axiomatic systems. We presented participants with a variety of “illusions”, which can be precisely defined mathematically, and asked them to adjust each “illusion” until they perceived it as dictated by mathematical principles. For example, where two geometrically parallel lines appear concave, as in the Hering illusion, subjects were asked to adjust the two lines until they appear perceptually parallel which, as our results show, entails a departure from the mathematical parallelism. Therefore, our overall conclusion is that mathematics does not consistently describe perceptual reality accurately.

Previous studies have shown that the magnitude of the perceptual illusory effect can be modified by adjusting various parameters (Knol et al., 2015; Restle & Decker, 1977). Here we go beyond by showing that whatever the degree of perceptual adjustment made to fit the mathematical axioms, the final result is irreconcilable with the mathematical one, in that achieving perceptually what the mathematical axiom suggests (e.g. that the two perceptually concave lines in the Hering illusion are in fact parallel) entails a significant departure from the mathematical axiom. We refer to these perceptual truths, as distinct from the mathematical ones, as *perceptual axioms* (Filippov & Zeki, 2022). We presume that these perceptual axioms are strong in that they rely on priors which are resistant to updating, even when cognitive knowledge of the mathematical reality is acquired.

### The Ebbinghaus illusion

We will use the Ebbinghaus illusion as an example with which to draw on the axioms of Euclidean geometry and present their irreconcilability with the perceptual axiom. Euclid’s eighth axiom states that “magnitudes that can be made to coincide are equal^”^ (Casey, 1885). Digitally and mentally the two inner disks in the Ebbinghaus illusion might be exactly superposed and be defined as equal according to this axiom. However, 96% of participants perceived inequality in this same state, and made an average adjustment of 37.4% of the initial disk diameter to reach a state in which the two disks are *perceived* as equal in size. These participants’ perceptual axioms, determined by their unique perceptual systems, were incompatible with the Euclidean axiom. The same is true for the other “illusions”. Although the magnitude of the adjustment may vary across studies (Knol et al., 2015; Mamassian & de Montalembert, 2010; Prinzmetal et al., 2001; Roberts et al., 2005), which can be explained by parameter differences (Knol et al., 2015; Restle & Decker, 1977), there nevertheless is a consistent discrepancy between perceptual and Euclidean truths.

### Incompleteness in mathematics

An early demonstration of incompleteness in mathematics came with the discovery of the sphere. Curved surfaces proved irreconcilable with certain axioms and propositions of Euclidean geometry, necessitating the formulation of a novel system under different axioms capable of dealing with curved surfaces, namely that of Riemannian geometry (Lee, 1997).

In the 1930s, David Hilbert and the formalist school of mathematics sought for a mathematical axiomatic system that was complete and consistent (Hilbert, 1930), one from which any statement formulated in the language of that system might be proven true or disproven. They also searched for an underlying set of axioms which were themselves proved to be consistent. In the first of his so-called *Incompleteness Theorems*, Kurt Gödel (1931) showed that within any system of first-order arithmetic logic there exist statements which are known to be syntactically true but can never be proven as such within that system. He also showed, in his second *Incompleteness Theorem*, that the consistency of the axioms of a logical system can never be proven within that system (Raatikainen, 2022). We take inspiration from Godel’s formulations to suggest that there may be more than one brain axiomatic system, and that the truths derived from one system may not be proven by truths derived from another logical system. Using the example of the Hering illusion given above, it cannot be proven within the perceptual axiomatic system, that the lines are indeed parallel, nor can it be shown within the Euclidean system that the lines may appear concave, suggesting that each axiomatic system is incomplete within itself. Thus, following on from Gödel, our results further expose incompleteness in reliance on any one logical system to obtain knowledge. The perceptual logical system has often been labelled “illusory” because it departs from what has been considered to be the “objective” and therefore truthful system derived from mathematics. We are not unaware of the further implications of this for knowledge derived from other brain logical systems. While others have discussed the existence of two different systems (e.g., Westheimer, 2008), we would like to emphasize that, in our view, both systems are axiomatic in the sense that their findings are accepted as true, but are irreconcilable; they follow separate logical systems in the brain that coexist but do not agree.

### Beyond the “ocular deception”

By labelling the stimuli used in this experiment as “illusions” psychologists and psychophysicists took a firm position – that these displays departed from the “objective” reality. Yet, the only reality that we are capable of experiencing is that which is generated by, and compliant to, brain mechanisms. The cognitively (mathematically) “objective” axioms of Euclidean geometry hold true within their system only; every effort made on a perceptual basis to make, for example, the two central disks in the Ebbinghaus illusion appear perceptually equal, will entail a departure from the objective mathematical condition. The same is true of other “illusions.”

Various pieces from the literature have utilised ideas from Bayesian probability as a conceptual basis for perception (Brown & Friston, 2012; Geisler & Kersten, 2002; Weiss et al., 2002; Yang et al., 2021; Zeki & Chén, 2020). Given that multiple environmental configurations can map onto the same sense data, this framework states that the brain must make inferences about the environment; these inferences are influenced by priors, which can be innate or learned. We suggest that the brain’s perceptual reality, which is exposed by optical “illusions”, relies on strong priors and is resistant to updating, even when cognitive knowledge of the mathematical truth is acquired. Although these studies propose frameworks to interpret these observations from a brain-based perspective, we hope that our work will inspire future studies to investigate the neural mechanisms involved in perceptual axioms and those derived from a more cognitive source.

The irreconcilable nature of geometric mathematical and perceptual axioms was implicitly recognised during the architectural and perceptual experiments of the great Greek architects (Ictinus and Callicrates) who designed the Parthenon. They deliberately modified the geometry of the massive Parthenon columns and spaced them more widely at the ends to counteract the visual effect of perceiving the columns as appearing unequal in size and in spacing when viewed from a distance (Notomi, 1999). The geometric modification was undertaken “To counteract the ocular deception by an adjustment of proportions” (Vitruvius, 20 B.C.E.). Here, Vitruvius, while implicitly acknowledging that perception trumps geometry, refers to the perceptual reality as a “deception”. In our experiments we proceeded in exactly the opposite direction — we counteracted the “geometric deception” by adjusting it to fit the “ocular reality” or, in our terms, to fit the perceptual axioms. By adjusting the presiding intuition around the truth of perceptual axioms such that it is recognised that perceptual truths are entirely compliant to brain mechanisms that generate logic and experience, we move beyond the stigma associated with so-called “optical illusions.” Even those mathematical truths considered real in a Platonic sense — that they are abstract forms that exist independently of human action – are still generated by brain mechanisms.

## CONCLUSION

The results presented here show that in the case of certain geometric pictures there exists a significant and irreconcilable difference between the perception of equality and its mathematical description. These pictures are commonly referred to as “optical illusions,” and are examples of cases in which perceptual truths are self-evident but cannot be proven by mathematics. Here we argue that these perceptual truths qualify as axioms and that the term “optical illusion” is a misnomer because it emphasises primarily the cognitive mathematical description of these pictures when in fact perception cannot be updated once this knowledge is acquired. Both perceptual and mathematical knowledge follow their own cogent logic, and in certain cases these systems are irreconcilable and reliant upon differing brain mechanisms. If mathematical sciences, including above all physics, aspires to a complete description of reality, such perceptual axioms must be imported into its canon.

## AUTHOR CONTRIBUTIONS

S.Z. conceived of the experiment; S.E.R. and A.B. programmed the stimuli used; Z.F.H. and S.E.R. ran the experiment; S.E.R., A.B., and Z.F.H. performed statistical analyses; Z.F.H., S.Z. and S.E.R. wrote the initial manuscript; all authors contributed to manuscript revisions.

## ACKNOWLEDGEMENTS

This work was supported by a Leverhulme Trust grant to Semir Zeki (RPG-2020-022). We thank M. Filippov for helpful discussion and review.

## DATA AVAILABILITY

The data that support the findings of this study are available from the corresponding author upon request.

## CODE AVAILABILITY

The code used in this study is available from the corresponding author upon request.

## ABBREVIATIONS

None.

## REFERENCES

Brainard, D. H. (1997). The Psychophysics Toolbox. Spatial Vision, 10(4), 433–436. 10.1163/156856897X00357

Brown, H., & Friston, K. J. (2012). Free-energy and illusions: The Cornsweet effect. Frontiers in Psychology, 3, 43. 10.3389/FPSYG.2012.00043/BIBTEX

Casey, J. (1885). The First Six Books of the Elements of Euclid (Third). Ponsonby and Weldrick.

Filippov, M., & Zeki, S. (2022). On the necessity of importing neurobiology into mathematics. PsyCh Journal. 10.1002/PCHJ.524

Geisler, W. S., & Kersten, D. (2002). Illusions, perception and Bayes. Nature Neuroscience, 5, 508–510. 10.1038/nn0602-508

Gödel, K. (1931). Über formal unentscheidbare Sätze der Principia Mathematica und verwandter Systeme I. Monatshefte Für Mathematik Und Physik, 38, 173–198. 10.1007/BF01700692/METRICS

Gray, J., & Ferreirós, J. (2022). Epistemology of Geometry. In E. N. Zalta (Ed.), The Stanford Encyclopedia of Philosophy (Fall ‘21). Metaphysics Research Lab, Stanford University.

Hilbert, D. (1930). David Hilbert’s Radio Address. https://www.maa.org/book/export/html/326610

Holt-Hansen, K. (1961). Hering’s Illusion. British Journal of Psychology, 52(4), 317–321. 10.1111/j.2044-8295.1961.tb00796.x

Knol, H., Huys, R., Sarrazin, J.-C., & Jirsa, V. K. (2015). Quantifying the Ebbinghaus figure effect: target size, context size, and target-context distance determine the presence and direction of the illusion. Frontiers in Psychology, 6, 1679. 10.3389/fpsyg.2015.01679

Lee, J. M. (1997). Riemannian Manifolds. Springer.

Mamassian, P., & de Montalembert, M. (2010). A simple model of the vertical–horizontal illusion. Vision Research, 50(10), 956–962. 10.1016/j.visres.2010.03.005

Notomi, N. (1999). The Unity of Plato’s Sophist: Between the Sophist and the Philosopher. Cambridge University Press. 10.1017/CBO9781107297968

Prinzmetal, W., Shimamura, A. P., & Mikolinski, M. (2001). The Ponzo illusion and the perception of orientation. Perception & Psychophysics, 63(1), 99–114. 10.3758/BF03200506

Raatikainen, P. (2022). Gödel’s Incompleteness Theorems. In E. N. Zalta (Ed.), The Stanford Encyclopedia of Philosophy (Spring ‘22). Metaphysics Research Lab, Stanford University.

Restle, F., & Decker, J. (1977). Size of the Mueller-Lyer illusion as a function of its dimensions: theory and data. Perception & Psychophysics, 21(6), 489–503. 10.3758/BF03198729

Roberts, B., Harris, M. G., & Yates, T. A. (2005). The roles of inducer size and distance in the Ebbinghaus illusion (Titchener circles). Perception, 34(7), 847–856. 10.1068/p5273

Vitruvius. (20 B.C.E.). De Architectura.

Weiss, Y., Simoncelli, E. P., & Adelson, E. H. (2002). Motion illusions as optimal percepts. Nature Neuroscience, 5, 598–604. 10.1038/nn0602-858

Westheimer, G. (2008). Illusions in the spatial sense of the eye: Geometrical–optical illusions and the neural representation of space. Vision research, 48(20), 2128–2142. 10.1016/j.visres.2008.05.016

Yang, S., Bill, J., Drugowitsch, J., & Gershman, S. J. (2021). Human visual motion perception shows hallmarks of Bayesian structural inference. Scientific Reports, 11, 3714. 10.1038/s41598-021-82175-7

Zeki, S., & Chén, O. Y. (2020). The Bayesian-Laplacian brain. European Journal of Neuroscience, 51(6), 1441–1462. 10.1111/ejn.14540

