## Supplementary Material for "Perceptual axioms are irreconcilable with Euclidean geometry"

#### SUPPLEMENTAL MATERIAL

Zachary F Hale [1], Samuel E Rasche [1], Ahmad Beyh [1], Semir Zeki [1,2]

[1] *Laboratory of Neurobiology, University College London, London, UK*
[2] *Corresponding author*

This document contains additional results related to the main experimental results reported in the paper.

### SCALING EFFECT

Preliminary studies involved presenting the five illusions discussed in the main manuscript to participants at three different scales on the screen, scaled by a factor of 0.5, 0.75, or 1. Scale factor 1 was used in the experiment proper. Subjects were asked to adjust the illusion until it appeared equal, and the adjustment relative to the initial size was recorded.

The results (Figure S1) confirmed that the size at which the illusions are presented does not significantly alter the extent of adjustment as a fraction of their initial size. There is no significant difference between the mean adjustment as a fraction of the initial size if the stimulus is presented at difference scales.

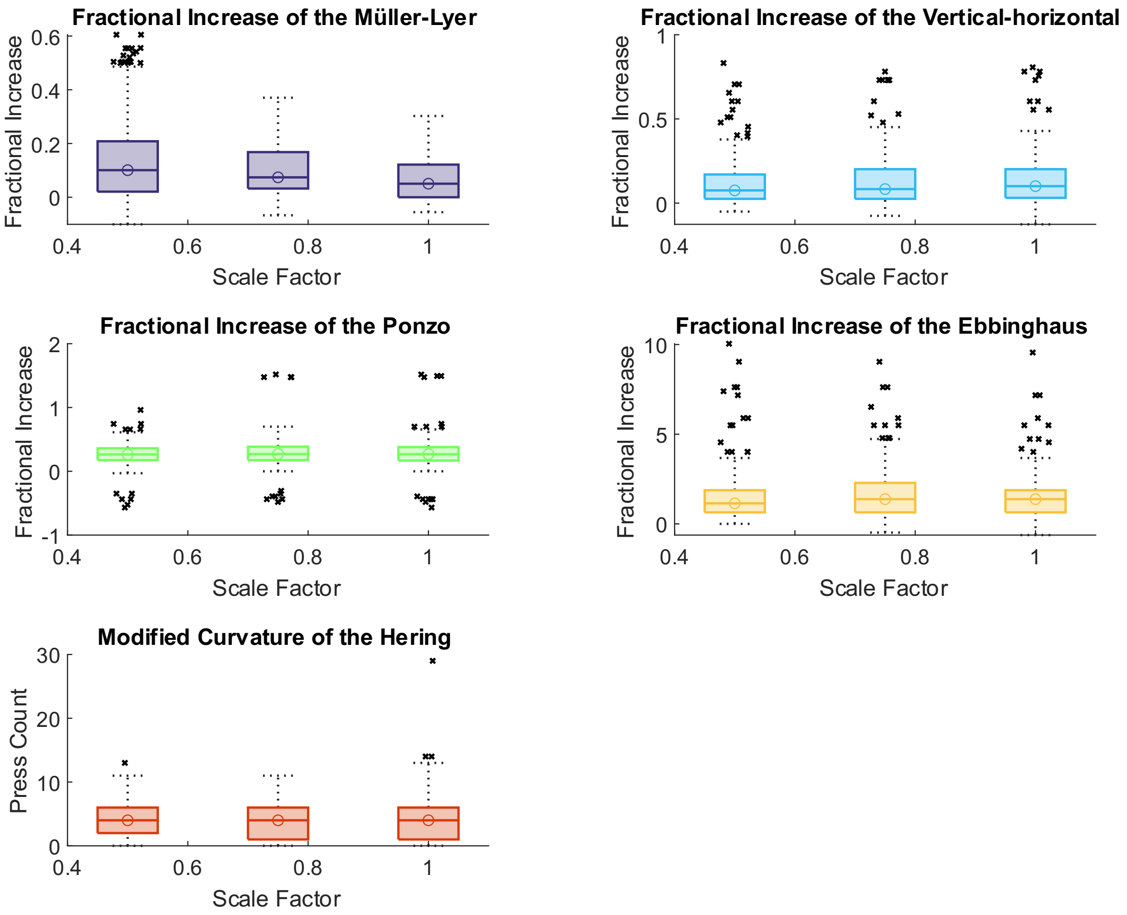


**Figure S1. Adjustment from mathematical to perceptual equality as a fraction of the initial length for each illusion.**
In this figure boxes span from the 25th to the 75th percentile of data and contain all instances of adjustment by 30 participants in the pilot phase. Centre lines and points indicate the median of each dataset. Some participants adjusted with a randomised increment, and others with a constant one.
